# The origin and diversification of pteropods predate past perturbations in the Earth’s carbon cycle

**DOI:** 10.1101/813386

**Authors:** Katja T.C.A. Peijnenburg, Arie W. Janssen, Deborah Wall-Palmer, Erica Goetze, Amy Maas, Jonathan A. Todd, Ferdinand Marlétaz

## Abstract

Pteropods are a group of planktonic gastropods that are widely regarded as biological indicators for assessing the impacts of ocean acidification (OA). Their thin aragonitic shells are highly sensitive to acute changes in ocean chemistry. However, to gain insight into their potential to adapt to current climate change, we need to accurately reconstruct their evolutionary history and assess their responses to past changes in Earth’s carbon cycle. Here, we resolve the phylogeny and timing of pteropod evolution with a phylogenomic dataset incorporating 21 new species and new fossil evidence. In agreement with traditional taxonomy, we recovered the first molecular support for a division between sea butterflies (Thecosomata: mucus-web feeders) and sea angels (Gymnosomata: active predators). Molecular dating demonstrated that these two lineages diverged in the early Cretaceous, and that all main pteropod clades, including shelled, partially-shelled and unshelled groups, diverged in the mid to late Cretaceous. Hence, these clades originated prior to and subsequently survived major global change events, including the Paleocene Eocene Thermal Maximum (PETM), which is the closest analogue to modern-day ocean acidification and warming. Our findings indicate that aragonitic calcifiers have been resilient to extreme perturbations in the Earth’s carbon cycle over evolutionary timescales.

## Introduction

Pteropods are marine gastropods that spend their entire life in the open water column. A remarkable example of adaptation to pelagic life, these mesmerizing animals have thin shells and a snail foot transformed into two wing-like structures that enable them to ‘fly’ through the water column (Fig. 1). Pteropods are a common component of marine zooplankton assemblages worldwide, where they serve important trophic roles in pelagic food webs, and are major contributors to carbon and carbonate fluxes in the open ocean (Berner and Honjo 1981; Hunt et al. 2008; Bednaršek et al. 2012a; Manno et al. 2018; Buitenhuis et al. 2019).

**Figure 1.**
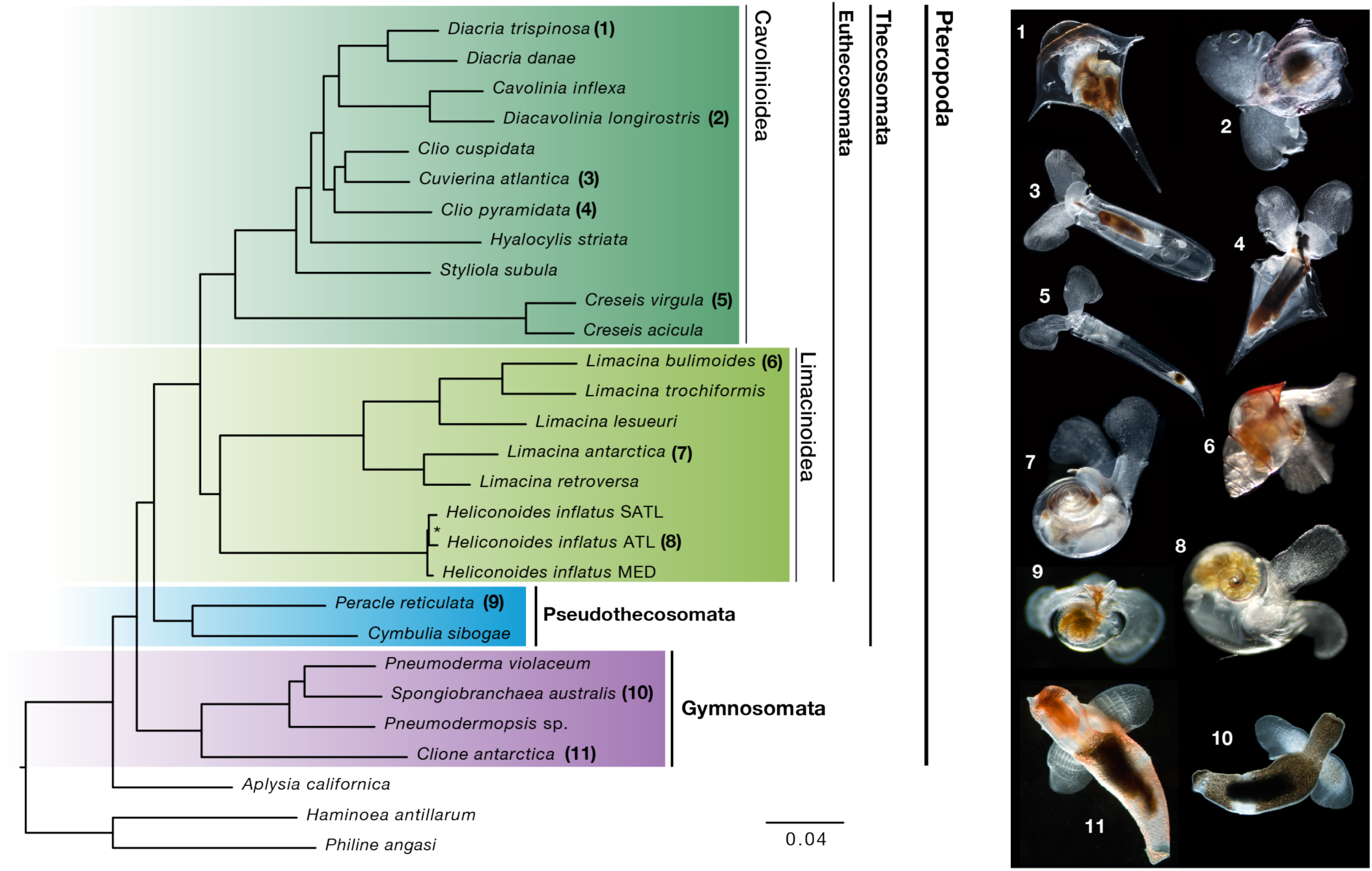
Phylogenomics resolves evolutionary relationships of pteropods. Euthecosomes (fully shelled species) and pseudothecosomes (ranging from fully shelled to unshelled species) are recovered as sister clades for the first time in a molecular analysis restoring the Thecosomata (‘sea butterflies’) as a natural group. Thecosomata and Gymnosomata (‘sea angels’) are monophyletic sister clades congruent with traditional morphology-based views. The superfamilies Cavolinioidea with uncoiled shells and Limacinoidea with coiled shells are also recovered as monophyletic sister clades. Maximum Likelihood phylogeny of 25 pteropod taxa, plus 3 outgroups assuming a LG+Γ model. The dataset comprises 2654 genes, concatenated as 834,394 amino acid positions with 35.8% missing data. A nearly identical topology is obtained modelled under CAT-GTR+Γ with a reduced dataset of 200 genes (Figure S1). Higher taxonomic divisions are indicated. All nodes receive maximal bootstrap support, except the node with an asterisk (bootstrap 95%). On the right are images of living pteropod species (numbers correspond with taxon labels, not to scale) collected and photographed by the authors. Scale bar indicates substitutions per site.

Shelled pteropods have been a particular focus for global change research because they make their shells of aragonite, a metastable form of calcium carbonate that is one and a half times more soluble than calcite (Mucci 1983; Sun et al. 2015). As their shells are susceptible to dissolution, pteropods have been called ‘canaries in the coal mine’, or sentinel species that signal the impacts of ocean acidification on marine calcifiers (e.g. Bednaršek et al. 2017a, Manno et al. 2017). Although shelled pteropods are already negatively affected in several regions of the global ocean (e.g. Bednaršek et al. 2012b; 2017b; Maas et al. in review), and will likely be seriously threatened if CO_2_ levels continue to rise (e.g. Moya et al. 2016; Maas et al. 2016; 2018), little is known about the evolutionary history of the group.

Improving the phylogenetic framework for pteropods and estimating the timing of divergence for major lineages will help determine what effect past periods of high atmospheric CO_2_, such as the Paleocene Eocene Thermal Maximum (PETM), have had on their diversification and survivorship. The PETM is widely regarded as the closest geological analogue to the modern rise in ocean-atmosphere CO_2_ levels, global warming and ocean acidification (Zachos et al. 2005; Hönisch et al. 2012; Penman et al. 2014). Knowing whether major pteropod lineages have been exposed during their evolutionary history to periods of high CO_2_ is important to extrapolate from current experimental and observational studies to predictions of species-level responses to global change over longer timescales.

Pteropods are uniquely suited to shed light on long-term marine evolutionary dynamics, because they are the only living metazoan plankton with a good fossil record (Bé & Gilmer 1977). The only other pelagic groups with abundant fossil records are protists, including foraminifers, radiolarians, coccolithophores, and extinct animal lineages, such as ammonites. The pteropod fossil record extends from a rare internal mold of *Heliconoides* sp. from the Campanian, late Cretaceous (72.1 Million years ago (Ma), Janssen & Goedert 2016) to the present, with abundant fossils from the Eocene onwards (from ∼ 56 Ma, reviewed in Janssen & Peijnenburg 2017). However, the fossil record of pteropods is somewhat limited because their shells are very thin and are only preserved in waters above the aragonite saturation depth, which is shallower than the saturation depth of calcite (Gerhardt & Henrich 2001). In addition, several groups of pteropods have only partial shells or are shell-less as adults and thus are rarely preserved in marine sediments. Hence, resolving the evolutionary history of pteropods requires a combination of molecular and fossil-based approaches to resolve past diversification and timing.

While most researchers recognise pteropods as being comprised of two orders, Thecosomata (‘sea butterflies’) and Gymnosomata (‘sea angels’), a recent taxonomic revision (Bouchet et al. 2017) identifies three suborders: 1) Euthecosomata, fully-shelled, omnivorous mucus-web feeders, 2) Pseudothecosomata, a poorly known group with shelled, partially-shelled and unshelled species that also use mucus webs for feeding, and 3) Gymnosomata, with shell-less adults that are specialised predators, primarily on euthecosomes. Progressive evolution towards loss of shells as an adaptation to planktonic life has been proposed for the group (Lalli & Gilmer 1989; Spoel & Dadon 1999), but never fully tested.

Previous attempts to resolve the molecular phylogeny of pteropods have relied on small subsets of genes, and resolution has been limited, especially at deeper nodes, due to large rate heterogeneity and insufficient taxonomic signal (Klussmann-Kolb & Dinapoli 2006; Jennings et al. 2010; Corse et al. 2013; Burridge et al. 2017). Here we used a phylogenomic approach based on transcriptome sequencing to fully resolve the phylogeny of pteropods. Using the pteropod fossil record to calibrate the timing of divergence, we estimate that two major groups of pteropods, ‘sea butterflies’ and ‘sea angels’ diverged in the early Cretaceous, and thus must have survived previous global perturbations to the ocean‘s carbonate system.

## Main text (results & discussion)

### Robust phylogenomic resolution

We generated new transcriptome data for 22 pteropod samples corresponding to 21 species collected along two basin-scale transects in the Atlantic Ocean (Table S1 and S2). We also incorporated available data for three additional pteropod species (Moya et al. 2016; Maas et al. 2018; Zapata et al. 2014) and three outgroup species: the sea hare *Aplysia californica*, within the proposed sister group of pteropods (Aplysiida), and two members of Cephalaspidea, *Haminoea antillarum* and *Philine angasi*, from Zapata et al. (2014). Our taxonomic sampling included representatives of all extant families of Euthecosomata, two of three families of Pseudothecosomata, and two of six families of Gymnosomata. All superfamilies of Pteropoda were sampled except Hydromyloidea (Gymnosomata).

We inferred a set of single-copy orthologues using gene family reconstruction from assembled and translated transcripts and selected 2654 single-copy nuclear genes for phylogenetic inference based on their taxonomic representation. These selected genes are well-represented in our transcriptomes with a median of 1815 genes per species, the least being 682 genes (77.4% missing data) for *Diacria trispinosa* (Table S2). We combined these single-copy orthologues in a large data matrix of 834,394 amino acids with 35.75% missing data.

Using the large data matrix (2654 genes) and a site-homogeneous model of evolution (LG+Γ_4_), we recovered a fully resolved phylogenetic tree with maximal support values at all interspecific nodes (Figure 1). To account for the putative limitations of site-homogeneous models that could lead to systematic error, we also applied a site-heterogeneous model (CAT+GTR+Γ) using Bayesian inference (Figure S1). For this analysis, we used a reduced data matrix comprised of the 200 most informative genes (108,008 amino acids, see Methods) and we recovered an identical topology but for a single terminal node (Figure S1).

### Pteropod systematics reappraised

Our trees confirm that pteropods are a monophyletic group with sea hares (*Aplysia*) as their closest sister group (Klussmann-Kolb & Dinapoli 2006; Zapata et al. 2014). Pteropods are split into two monophyletic clades: Thecosomata and Gymnosomata, which is congruent with the traditional classification but strongly supported by molecular evidence for the first time. An obvious shared character for pteropods are the wing-like structures or ‘parapodia’ used for swimming. Histological and ultrastructural studies showed that the muscle arrangements in parapodia are very complex and look similar in Thecosomata and Gymnosomata, supporting a homologous origin (Klussmann-Kolb & Dinapoli 2006). Thecosomata comprise the Euthecosomata and Pseudothecosomata clades, whose member species are all omnivorous mucus-web feeders (Gilmer & Harbison 1986). Their common feeding mechanism is reflected in the well-developed mucus secreting pallial gland that is shared among all thecosomes, as well as a muscular gizzard with which they can crush the hard exoskeletons of their prey (Lalli & Gilmer 1989). Gymnosomata are shell-less at the adult stage and are specialized carnivorous hunters. They have several morphological characters that set them apart from Thecosomata, including tentacle-like structures called ‘buccal cones’ and hook sacs to grab and manipulate shelled pteropod prey (Lalli 1970, Klussmann-Kolb & Dinapoli 2006). Hence, the recent revision by Bouchet et al. (2017) with three separate suborders (Euthecosomata, Pseudothecosomata, Gymnosomata) should revert back to the original classification with two main clades: Thecosomata, comprising the Euthecosomata and Pseudothecosomata, and Gymnosomata as shown in Figure 1. Within the fully-shelled Euthecosomata, we obtained maximal support for the coiled (Limacinoidea) and uncoiled (Cavolinioidea) superfamilies as monophyletic sister clades for the first time in a molecular analysis. These results finally stabilize the higher-level taxonomy of the group, which has been debated ever since Cuvier (1804) established the Pteropoda as a separate order of molluscs.

In agreement with previous molecular phylogenetic analyses (Corse et al. 2013; Burridge et al. 2017), we find good support for lower level groupings (e.g. genera *Diacria* and *Limacina*) and recover *Creseis* as the earliest diverging lineage within the uncoiled shelled pteropods. The genus *Clio*, however, is paraphyletic in our analyses with *Clio cuspidata* and *Cuvierina atlantica* grouping together, and *Clio pyramidata* either as a sister taxon to this group (maximum-likelihood tree, Fig. 1) or to the clade *Cavolinia* + *Diacavolinia* + *Diacria* (Bayesian tree, Fig. S1). These results are congruent with the branching obtained using a wider taxon sampling but only three genes (Burridge et al. 2017). Thus, it seems plausible that the genus *Clio* consists of two distinct groups, which could be characterised by distinct larval shell shapes, including *C. cuspidata* and *C. recurva* in one clade, and *C. pyramidata* and *C. convexa* in the other (Janssen 2012). Sampling of additional species for transcriptome sequencing and more detailed morphological analysis are necessary to definitively revise the taxonomy of this genus.

### Divergence times of major pteropod lineages

Estimating divergence times based on genome-scale datasets has been shown to be accurate and powerful, however, this depends on the use of realistic evolutionary and clock models as well as a reliable fossil calibration scheme (Tanner et al. 2017; Irisarri et al. 2017). We inferred divergence times of the main pteropod lineages using a recent revision of their fossil record (Janssen & Peijnenburg 2017), which provided eight meaningful calibrations (Fig. 2, Table S3). Within the sister clade of Pteropoda (Aplysiida) the shelled genus *Akera* has by far the best and oldest known fossil record (Cossmann 1895a, 1895b, Valdes & Lozouet 2000) and we chose the oldest confidently identified species of *Akera* (*A. neocomiensis*) to provide the calibration for the Aplysiida + Pteropoda clade (Table S3). To ensure accurate reconstructions, we chose a realistic model of sequence evolution (CAT-GTR) using the reduced data matrix of 200 genes. We performed cross-validation to select the best clock model (CIR, Figure 2) but also verified that the divergence times reported under alternative clock models, such as autocorrelated log-normal or uncorrelated gamma multipliers, were not markedly different for the major clades (Figure S2, Table S4).

**Figure 2.**
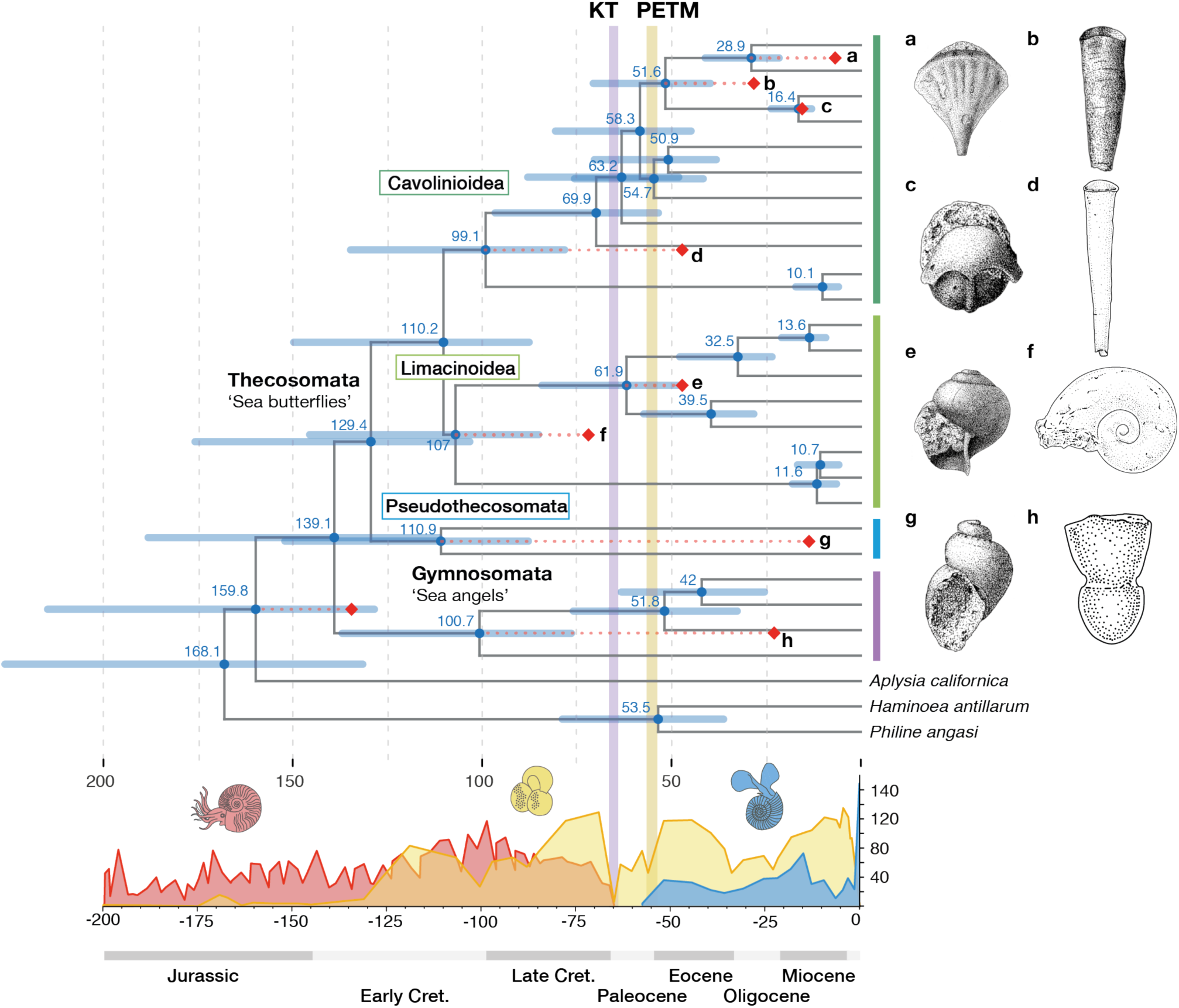
All major lineages of pteropods diverged in the Cretaceous and thus must have survived previous major global change events. Chronogram of pteropods (25 taxa and 3 outgroups) based on 200 genes, concatenated as 108,008 amino acid positions, analysed under a CAT-GTR+Γ evolutionary model and CIR relaxed clock model. Red diamonds indicate fossil-calibrated nodes (see Table S3) with red dotted lines to show divergence from minimum ages of the fossils. Two major global change events are indicated as KT (Cretaceous-Tertiary asteroid impact, 66 Ma) and PETM (Paleocene Eocene Thermal Maximum, ∼56 Ma). Curves for ammonite (red), planktonic foraminifer (yellow), and pteropod (blue) species diversity through time are based on sediment records from Yacobucci (2015), Boudagher-Fadel (2015), and our own data base (available from the authors on request), respectively. Drawings of fossil species used for calibrations correspond to nodes a: *Diacria sangiorgii* (7.2 Ma), b: *Vaginella gaasensis* (28.1 Ma), c: *Cavolinia microbesitas* (16 Ma), d: *Euchilotheca ganensis* (47.8 Ma), e: *Limacina gormani* (47.8 Ma), f: *Heliconoides sp*. (72.1 Ma), g: *Peracle amberae* (16 Ma), and h: *Clione ? imdinaensis* (23 Ma). The oldest outgroup calibration is *Akera neocomiensis* (133 Ma, no drawing available).

Our phylogenomic time-tree dates the origin of pteropods to the early Cretaceous (139.1 Ma with 95% credibility interval (CrI): 188.2-111.4, Fig. 2), which is substantially older than the estimates based on previous molecular phylogenies (Paleocene and late Cretaceous as estimated by Corse et al. 2013 and Burridge et al. 2017, respectively). While previous estimates were based on a few markers, our analyses are based on a much more comprehensive gene set and rely on more realistic models of evolution and better-curated fossil calibrations. We also find that divergences of the two main lineages, Thecosomata and Gymnosomata, are placed in the Cretaceous period, dated at 129.4 (CrI: 175.8-103.3) and 100.7 (137.0-76.4) Ma, respectively. Of the fully shelled pteropods, all coiled species share a most recent common ancestor at 107 (145.5-85.1) Ma and the uncoiled species at 99.1 (134.7-78.2) Ma.

Different groups of pteropods (sea butterflies and sea angels, species with coiled and uncoiled shells) were already present in the Cretaceous and thus must have survived previous major environmental changes and associated extinctions such as the asteroid impact at the end of the Cretaceous (KT, 66 Ma, Smit & Hertogen 1980; Schulte et al. 2010) and the Paleocene Eocene Thermal Maximum (PETM, 56 Ma, Kennett & Stott 1991; Dickens et al. 1997; Zachos et al. 2005) (Fig. 2). Many pelagic groups disappeared during the well-known mass extinction at the KT boundary (Jablonski 1994), but the multiple environmental factors that changed simultaneously during this period make it difficult to pinpoint a specific cause of these changes, such as ocean acidification (Hönisch et al. 2012). The PETM had an impact on marine calcifiers as shown by a rapid decrease in calcium carbonate content in marine sediments (Luo et al. 2016) and the recording of one of the largest extinctions among deep-sea benthic foraminifers (Kennett & Stott 1991; Schmidt et al. 2018). Planktonic calcifiers were also affected during the PETM, with shifts in species assemblages and changes in species abundance of calcareous nannoplankton, but no clear signal of increased extinction (Gibbs et al. 2006, Bown & Pearson 2009; Kelly et al. 1996; Raffi et al. 2009). For the pteropod fossil record, we see an increase of species from pre-PETM (Thanetian, 2 species) to post-PETM (Ypresian, 32 species) which led to the suggestion that the PETM might have had a triggering effect on pteropod evolution (Janssen et al. 2016).

### Evolutionary history of pteropods

Our estimated divergence times for the major pteropod groups predate by far the oldest known fossils for these groups (Fig. 2). The largest discrepancies are found for Gymnosomata and Pseudothecosomata, for which the oldest fossils are from the Chattian (probably a *Clione* sp., 28-23 Ma) and Chattian-Burdigalian periods (*Peracle amberae*, 28-16 Ma), respectively (Janssen 2012). This discrepancy is not surprising as these groups are characterised by reduced shells or no shells at all as adults and their microscopic (larval) shells remain mostly uncharacterised. The pteropod fossil record is generally affected by a strong taphonomic bias as their delicate aragonitic shells preserve poorly. Furthermore, micropaleontologists have traditionally focused on calcitic planktonic calcifiers (coccolithophores, foraminifers) rather than aragonite-producing ones, such as pteropods. This trend is illustrated by plotting the reported diversity of pteropod species through time (Figure 2), showing a sharp increase in the number of recent species (162) compared to previous periods (21 species for the Pleistocene and a median of 34.5 species for the preceding Neogene periods). Since pteropods are considered to be promising new proxy carriers in palaeoceanography, recording surface ocean temperature and carbonate ion concentrations (Wall-Palmer et al. 2012; Keul et al. 2017), this renewed interest will hopefully result in a reappraisal of their fossil record in the coming years.

Surprisingly, we find similar Cretaceous origins for the coiled (Limacinoidea) and uncoiled (Cavolinioidea) shelled pteropods even though the fossil record for coiled shells extends much further (Campanian, late Cretaceous) than for uncoiled shells (Ypresian, early Eocene). The species with straight, bilaterally symmetrical shells were thought thought to have derived from coiled ancestors (*Altaspiratella*) that show a trend of despiralisation in the fossil record starting during the early Ypresian and giving rise to the oldest representatives of Cavolinioidea (*Camptoceratops* and *Euchilotheca*) (first suggested by Boas (1886) and reviewed in Janssen & Peijnenburg 2017). It is interesting to note that shell microstructure differs fundamentally between coiled and uncoiled pteropods. Uncoiled species have an outer prismatic layer and a thick inner layer with a unique helical microstructure (Bé et al. 1972; Li et al. 2015), whereas coiled shells have simple prismatic and crossed lamellar microstructures (Lalli & Gilmer 1989), similar to the outgroup *Aplysia* (Marin et al. 2018). Examination of shell microstructures of fossil species could shed further light on the phylogenetic placement of the fossils. For instance, the fossil *Camptoceratops priscum* from the early Eocene was found to have helical microstructure similar to extant Cavolinioidea (Curry & Rampal 1979). A recent report by Garvie et al. (in review) of potentially much older uncoiled pteropod-like fossils from Mesozoic and Paleocene rocks of the southern United States found that their microstructure is, in contrast, crossed-lamellar. If future analyses show that these are indeed pteropods, despiralisation must have occurred multiple times throughout their evolutionary history. Reports of Cretaceous pteropods remain extremely rare despite considerable paleontological effort, which suggests that pteropods were not abundant and/or very poorly preserved during this period. Notably, Cretaceous deposits are often dominated by limestones, which are unsuitable for the preservation of thin-walled aragonitic pteropod shells (Janssen & Peijnenburg 2017).

We found that the shelled groups (sea butterflies) diverged before the unshelled ones (sea angels), suggesting that unshelled species evolved from shelled ancestors (Fig. 2). However, we do not see a clear trend toward gradual loss of shells in our phylogeny. This could be further assessed by sampling more species belonging to the elusive Pseudothecosomata, with species ranging from fully-shelled (*Peracle* spp.) to partially-shelled (*Cymbulia, Corolla, Gleba* spp.) and even entirely unshelled as adults (*Desmopterus* spp.). In our phylogeny, we only included one *Peracle* species possessing an external coiled calcareous shell and one *Cymbulia* species bearing a gelatinous ‘pseudoconch’. Since the pteropod outgroups are benthic gastropods (Aplysiida, Cephalaspidea) with mostly coiled or reduced shells, the ancestor of pteropods most likely lived on the seafloor and had a coiled shell (Klussmann-Kolb & Dinapoli 2006; Corse et al. 2013). Some authors proposed that pteropods evolved in a neotenic fashion, where larvae of benthic gastropods became sexually mature and lived their full life cycle in the open water column, because of ‘juvenile’ shell characters such as the sinistral spiral and aragonitic shell structure (Lemche 1948; Huber 1993). However, this hypothesis has been questioned by Jägersten (1972), who argued that the foot – an adult feature – played a decisive role in the transition from a benthic to a holoplanktonic existence by transforming from a creeping organ to swimming fins. It is interesting to note that a similar hypothesis was proposed for the only other extant group of holoplanktonic gastropods, the heteropods (Pterotrachoidea). For this group, the earliest fossil, *Coelodiscus minutus*, dates back to the early Jurassic and represents the oldest known holoplanktonic gastropod (Teichert & Nützel 2015). Colonization of the open water column required numerous adaptations in both groups of holoplanktonic gastropods independently: pteropods belonging to Heterobranchia and heteropods belonging to Caenogastropoda.

### Fate of sea butterflies and angels from Cretaceous to Anthropocene

Pteropods evolved in the early Cretaceous and thus were contemporaries of other major calcifying groups in the open ocean, such as ammonites and foraminifers (Fig. 2). Based on their fossil records, Tajika et al. (2018) suggested that planktonic gastropods filled the ecological niche left empty by juvenile ammonites after their extinction at the end of the Cretaceous. However, this seems less likely given our earlier estimated origins of both sea butterflies (thecosomes) and sea angels (gymnosomes). Instead, we propose that sea angels evolved as specialized predators of sea butterflies during the Cretaceous, perhaps in an evolutionary arms race (Seibel et al. 2007) in which thecosomes evolved stronger and ever more sophisticated shells (e.g. *Diacria, Cavolinia*) and gymnosomes evolved adaptations for prey capture and extraction, specific to the shell shapes of their prey. For instance, *Clione* feed exclusively on certain *Limacina* species by manipulating the coiled shells with their flexible buccal cones and extracting the prey with their hooks, whilst *Pneumodermopsis* feed on long and straight-shelled *Creseis* species using their long and flexible proboscis for prey extraction. There are also accounts of unsuccessful attacks, which happen when sea butterflies retract rapidly enough and far enough into their shell (Conover & Lalli 1972, Lalli & Gilmer 1989). Moreover, even though thecosomes are eaten by a variety of predators ranging from planktonic crustaceans to fish, whales and seabirds, no other pelagic group than gymnosomes feeds predominantly on thecosomes. Hence, we consider it plausible that different groups of pteropods co-evolved during a period that is referred to as the Marine Mesozoic Revolution. In this major evolutionary episode, a series of ecological shifts took place on the sea floor because of the evolution of powerful, relatively small, shell-destroying predators, including a massive radiation of predatory gastropods in the Early Cretaceous (Vermeij 1977; Harper 2003). This forced benthic gastropods to develop heavily armoured shells and perhaps also to escape into the open water column, as was suggested for earlier geological periods (Signor & Vermeij 1994, Teichert & Nützel 2015). Furthermore, during the mid-late Cretaceous major changes in ocean circulation, stratification and nutrient partitioning took place which were favorable for plankton evolution, in particular for planktonic calcifiers (Leckie et al. 2002).

Although the open ocean may have been a refuge for gastropods in the early Cretaceous, it is an increasingly challenging habitat to survive in during the Anthropocene. Planktonic gastropods have evolved thin, fragile shells of aragonite that are sensitive to ocean acidification, and most species live in surface waters where CO_2_ is absorbed by the ocean. Incubation experiments with shelled pteropods mimicking future ocean conditions have shown that elevated pCO_2_ and undersaturated aragonite conditions cause decreased calcification rates (Comeau et al. 2012), shell dissolution (Lischka et al. 2011), increased mortality (Bednaršek et al. 2017b; Thabet et al. 2015) and differential expression of genes involved in neurofunction, ion transport, and shell formation (Moya et al. 2016; Maas et al. 2018). However, such experiments have primarily assessed phenotypic responses in short-term, single-generation studies, and thus cannot take into account the abilities of organisms to acclimate or adapt to changing conditions over longer timescales. The geological record, alternatively, can provide insight into long-term evidence of ocean acidification and the associated responses of marine calcifiers. However, the fossil record of pteropods appears far from complete and hence, we need to rely on estimates of molecular divergence times to resolve the tempo and pattern of their evolution. The fact that pteropods have survived previous episodes of ocean acidification, such as during the PETM, does not, however, mean that they are infinitely resilient to current changes. Current rates of carbon release from anthropogenic sources are at least an order of magnitude higher than we have seen for the past 66 million years (Zeebe et al. 2016). Although our results suggest resilience of pteropods to past ocean acidification, it is unlikely that they have ever, during their entire evolutionary history, experienced global change of the magnitude and speed that we see today.

## Acknowledgements

We appreciate the comments of J. Paps, D. Gavriouchkina and M. Malinsky on earlier versions of this manuscript. We acknowledge the Oxford Genomics Centre, Wellcome Trust Centre for Human Genetics, for performing the transcriptome sequencing. We would like to thank A.K. Burridge for collecting samples and T. Smythe, G. Tarran, and A. Rees for cruise leadership and support for participation in the AMT programme. This study contributes to the international IMBeR project and is contribution number 339 of the AMT programme.

## Author contributions

KTCAP conceived this study and obtained funding. KTCAP and EG collected and identified specimens at sea. KTCAP and FM carried out the labwork. FM led the data analyses. KTCAP and AWJ reviewed the fossil record and provided calibrations. KTCAP and FM drafted the manuscript, and all others commented.

## Methods

### Sample collection

Pteropod specimens were collected on Atlantic Meridional Transect (AMT) cruises 22 and 24 (2012, 2014), using 0.71m diameter bongo and RMT1 nets towed obliquely between a median of 305 m depth and the sea surface (200, 333 µm nets; Table S1). Animals were sorted and identified live from bulk plankton, preserved in RNALater (Invitrogen) and flash frozen in liquid nitrogen, with storage at −80°C.

### Transcriptome sequencing and filtering

RNA was extracted using RNAeasy micro or mini kits (Qiagen) after homogenisation with a Tissuelyser (Qiagen). The number of individuals used for extraction varied from 1, in most cases, to 10 (Table S1). RNA quantity was determined by fluorimetry using a Qubit (Invitrogen) and RNA integrity was assessed using the Xperion system (Biorad). RNA-seq libraries were prepared using the TruSeq RNA Library preparation kit (Illumina) and between 12 and 33M paired-end reads per sample were sequenced for 100 cycles on a HiSeq2000 platform at the Wellcome Trust Centre for Human Genetics (Oxford). 2.3 to 6.6 Gb of DNA sequence data were obtained from each of 22 samples, ensuring accurate *de novo* transcriptome assemblies (Table S2). Reads were deposited to the SRA under the bioproject accession XXXX. After quality assessment with Fastqc (Andrews 2010), reads were trimmed using sickle (Joshi 2011) and subsequently assembled using *Trinity* (v2.3.2) with default parameters and a k-mer of 25 (Grabherr et al. 2011).

To avoid cross-contamination between samples sequenced on the same Illumina lanes (possibly due to index ‘hopping’), we applied a filtering procedure based on relative transcript expression across datasets, similar to the one implemented in Croco (Simion et al. 2018). Briefly, for the assembled transcriptome of each dataset, we measured expression level using Kallisto (v0.42.4) (Bray et al. 2016) in each of the multiple datasets sequenced simultaneously. We calculated the ratio of TPMs between the putatively contaminating datasets and the original dataset, and excluded transcripts with a contaminant enrichment greater than two-fold enrichment, and a minimal count lower than 2 in the original dataset. Across datasets, a median 6% of transcripts were excluded on the first criterion and a median 26% on the second.

*De novo* assembled transcriptomes usually include a high degree of redundancy, as alternative transcripts derived from the same genes are distinguished in the assembly. As this might constitute a problem for orthology assignment, we clustered transcripts based on the fraction of remapped reads that they share. To do so, we mapped reads from transcriptomes back to transcripts using Bowtie2 enabling up to 50 multi-mappers (-k 50) and processed the resulting alignments with Corset (v1.06) (Davidson and Oshlack 2014). Then, we estimated transcript expression using Kallisto (v0.43.1) (Bray et al. 2016) and we selected the most highly expressed transcript with each Corset cluster as a reference transcript for subsequent steps. The best open reading frame (ORF) for each of these selected transcripts was predicted using Trans-decoder (v5.0.2) using a blast against a version of swissprot limited to metazoan taxa (e-value 10^−5^) (Haas et al. 2008).

### Phylogenetic analyses

We conducted orthology inference using the OMA package (v2.3.0), which performs smith-waterman alignment and identifies orthologues based on evolutionary distance (Roth et al. 2008). We selected single-copy orthologues (OMA groups) represented by at least one of the three outgroup taxa (*Aplysia californica, Haminoea antillarum* or *Philine angasi*) and at least half of the 28 ingroup taxa. These cut-offs yielded 2654 single-copy orthologues suitable for phylogenetic analysis. For each orthologue family, protein sequences were aligned using Mafft and poorly aligned regions trimmed using Trimal (Capella-Gutiérrez et al. 2009) for sites with a gap in more than 75% of taxa (-gt 0.15) and applying a low minimum similarity threshold (-st 0.001). The concatenation of these 2654 orthologuous genes yields a supermatrix of 834,394 positions with a total fraction of 35.75% missing data. We performed Maximum-likelihood reconstruction with ExaML (v3.0.17) using a partitioned site-homogeneous model (one partition per orthologue) and a LG+Γ_4_ model (Kozlov et al. 2015). Node support was calculated using 100 bootstrap replicates inferred with independently generated starting trees (Figure 1). To compare results with a site-heterogeneous model, we used Bayesian inference and a reduced dataset (because the complete dataset would be computationally intractable). We selected a subset of 200 genes which showed the maximal average bootstrap support when analysed independently (using RAxML v8.1.18, LG+Γ_4_ model and 100 rapid bootstraps) (Stamatakis 2014). The concatenation of these 200 genes generated a 108,008 amino acid alignment. This alignment was analysed using Phylobayes-MPI (v1.6) assuming a CAT+GTR+Γ_4_ model) with chains run for more than 1500 generations with 500 discarded as burnin. Chain convergence was checked and maxdiff found at 0 (Rodrigue & Lartillot 2014).

### Molecular divergence time inference

We used Phylobayes (v4.1c) to infer molecular divergence times using the reduced 200 gene supermatrix and the topology from Figure 1 (ML analysis) (Lartillot et al. 2009). We used the CAT+GTR+Γ_4_ model of sequence evolution and we assayed several clock models, the lognormal autocorrelated process (-ln), the CIR process and the uncorrelated gamma multiplier process, with a birth-death prior on divergence time, and soft bounds on calibration points (Lepage et al. 2007). We applied 9 calibration points with uniform priors based on the fossil records of pteropods (Table S3) and a root prior of 150 Myr with a standard deviation of 70 Myr. Using a 10-fold cross validation procedure, we found a best fit of the CIR process over log-normal and uncorrelated gamma processes (Figure 2 and Figure S2). The root prior was conservatively chosen based on the oldest known heterobranch (e.g. *Cylindrullina* dated 240 Ma, (Ponder & Lindberg 2008)) and *Akera* fossils (*Akera mediojurensis* and *Akera neocomiensis* dated at 163-166 Ma and 133 Ma, respectively, Cossmann 1895a;b; Cossmann 1896). Broad deviations were used to account for the uncertainty of the fossil record. The consistency of the root prior was assessed by running chains without the data, which yielded a root age of 154 ± 32 Myr.

## Data availability

Alignments and files generated during analyses including Bayesian sample and bootstrap replicates are available at the following address: XXXX.

## Supplementary Figures

**Figure S1.**
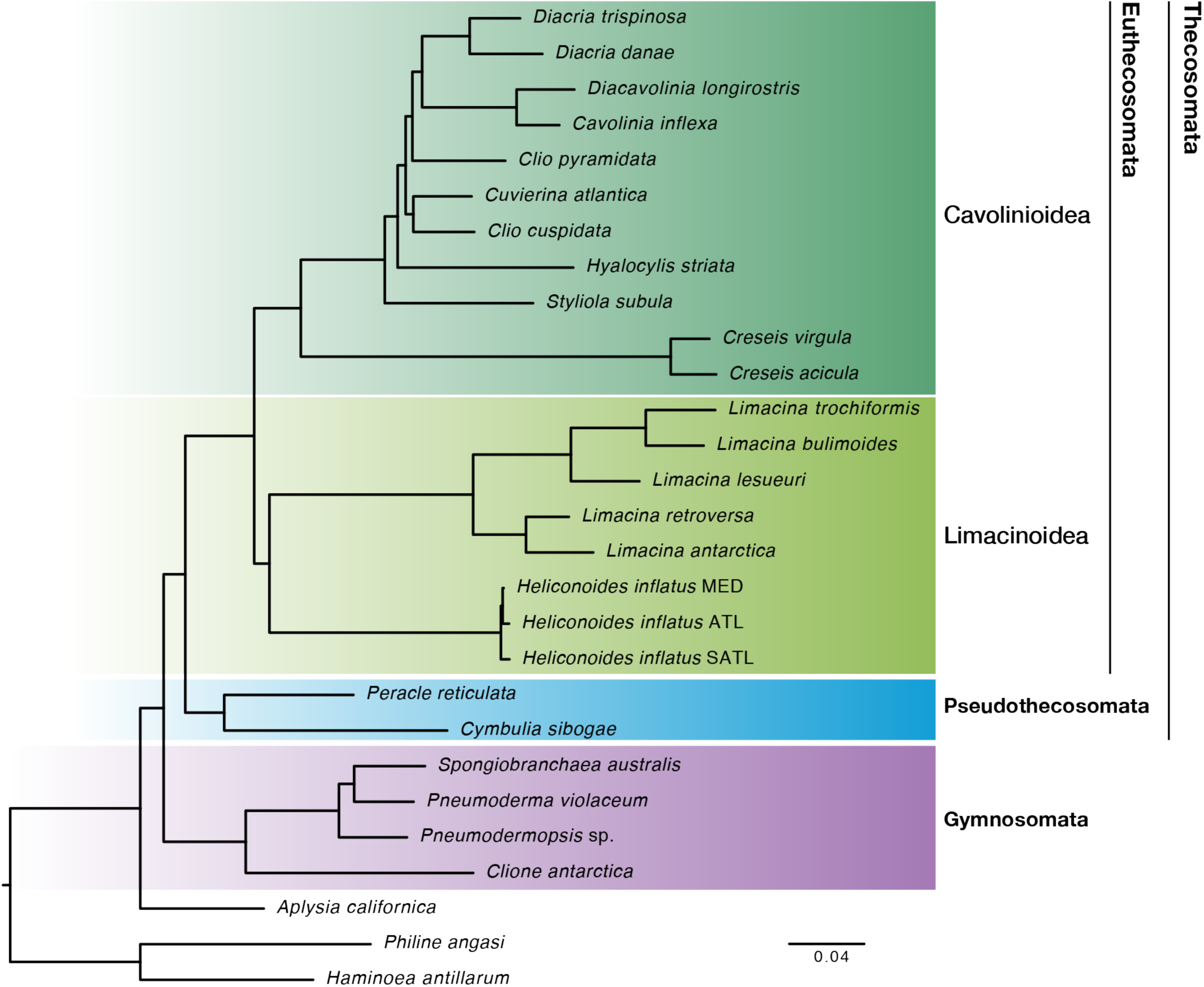
Phylogenomics resolves evolutionary relationships of pteropods. Euthecosomes (fully shelled species, green) and pseudothecosomes (ranging from fully shelled to unshelled species, blue) are recovered as sister clades for the first time in a molecular analysis restoring the Thecosomata (‘sea butterflies’) as a natural group. Thecosomata and Gymnosomata (‘sea angels’, purple) are monophyletic sister clades congruent with traditional morphology-based views. The superfamilies Cavolinioidea with uncoiled shells and Limacinoidea with coiled shells are also recovered as monophyletic sister clades. Bayesian phylogeny of 25 pteropod taxa, plus 3 outgroups assuming a CAT+GTR+Γ_4_ model using Phylobayes-MPI. The dataset was comprised of 200 selected genes (see Methods), concatenated as 108,008 amino acid positions. All nodes received maximal posterior probabilies.

**Figure S2.**
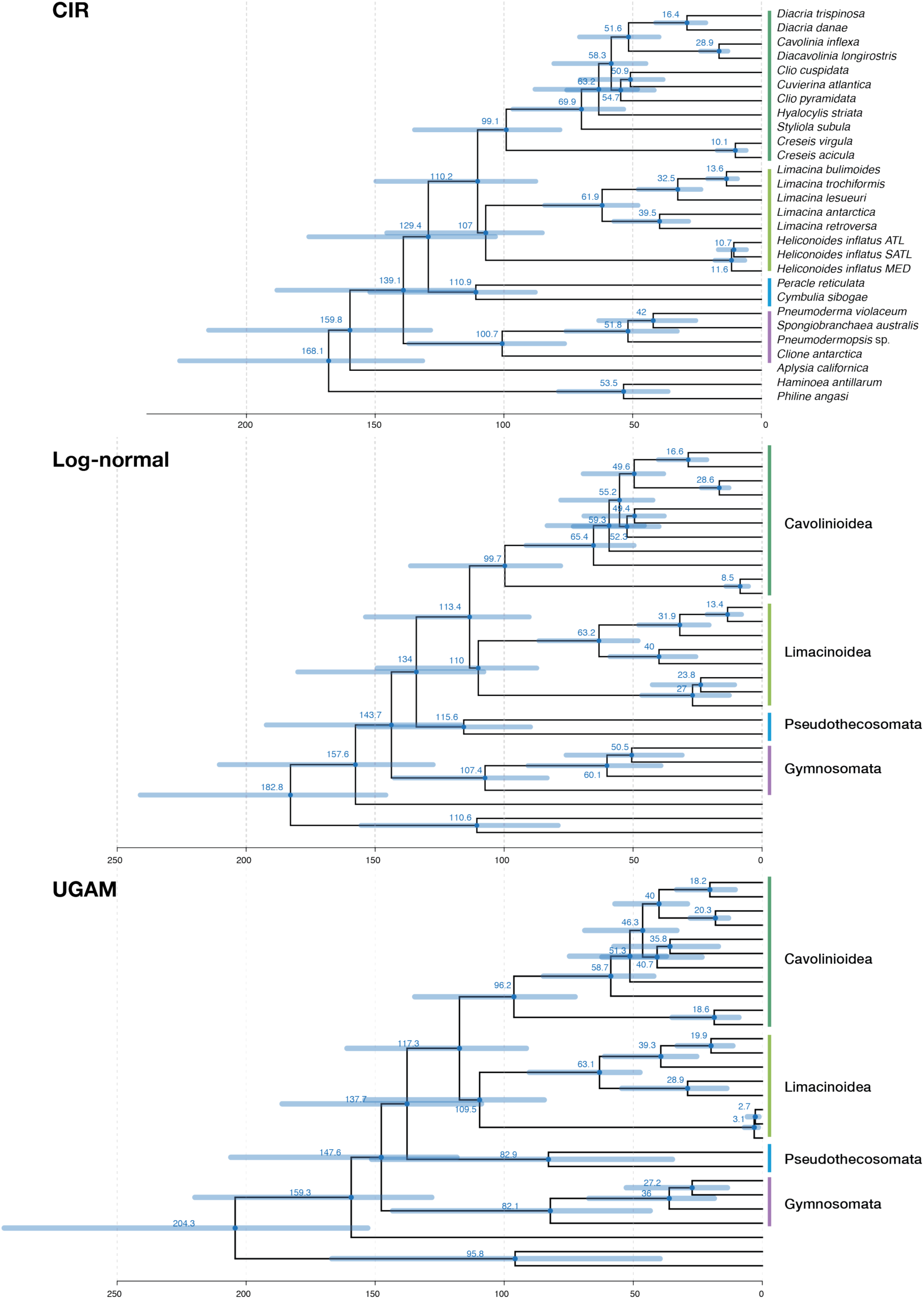
Divergence times for major pteropod clades under different clock models were not markedly different. Bayesian phylogenies to infer molecular divergence times using the reduced 200 genes supermatrix, the topology of Figure 1 and a CAT+GTR+Γ_4_ model of sequence evolution (see Methods) with three different clock models: the CIR process (CIR, see also Figure 2), the lognormal autocorrelated process (Log-normal), and the uncorrelated gamma multiplier process (UGAM).

## Supplementary tables

**Table S1.**
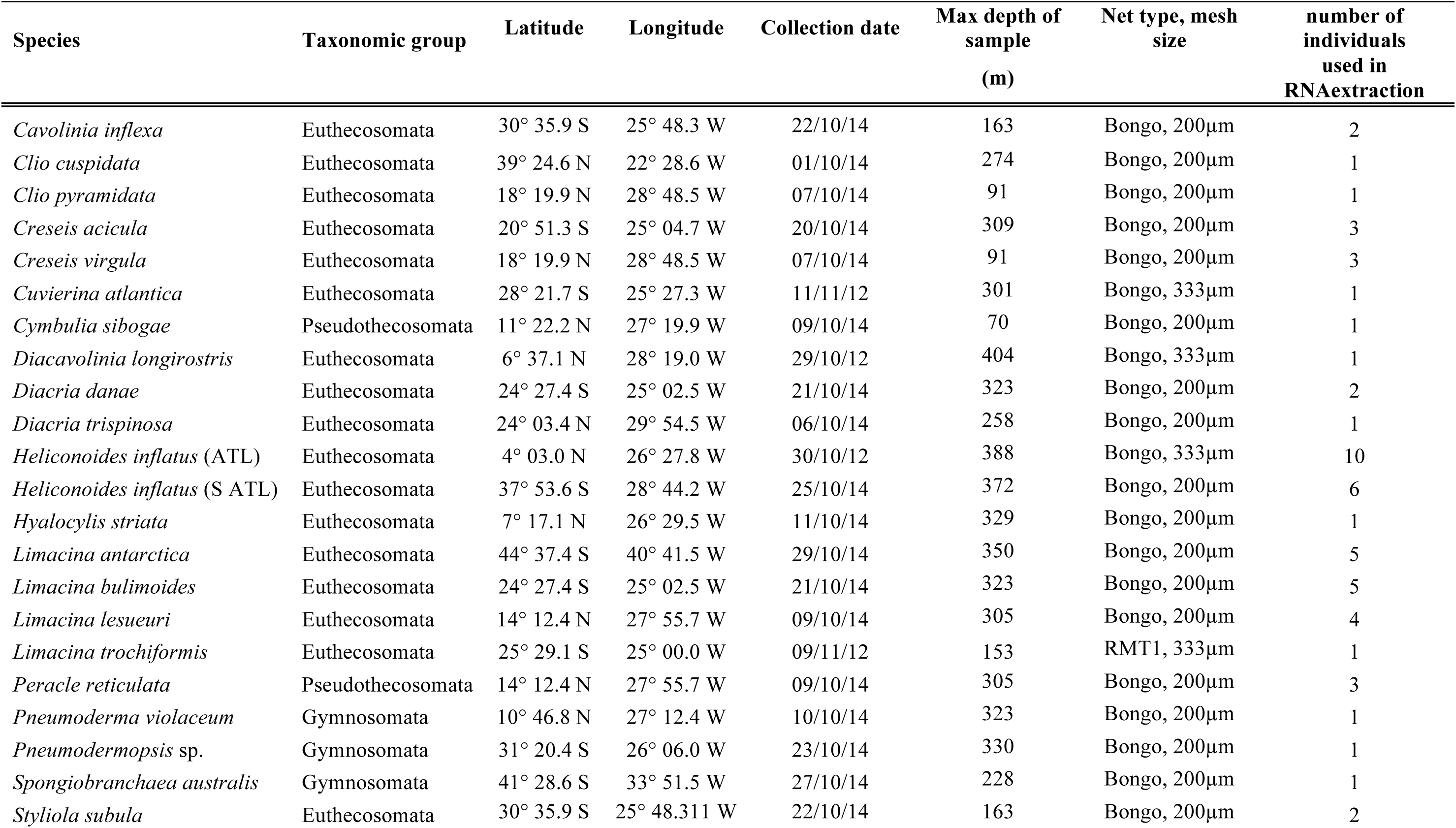
Sampling details of pteropod taxa for which new transcriptome data were collected.

**Table S2.** Details of 28 transcriptome datasets used in phylogenomic analyses. N_reads is the number of sequenced paired-end reads, N_pept is the number of predicted peptides among transcripts. N_OG is the number of selected single-copy orthologues recovered for this species. P_miss_OG is the percentage of missing single-copy orthologues for the taxon, P_gaps is the fraction of gaps and P_miss_tot is the total fraction of missing data in the alignment for the taxon (gaps and missing genes).

| Species | Order | Taxonomic group | accession | N_reads | N_pept | N_OG | P_miss_OG | P_gaps | P_miss_tot |
| --- | --- | --- | --- | --- | --- | --- | --- | --- | --- |
| *Aplysia californica* | Aplysiida | Outgroup | ftp.broadinstitute | - | 92469 | 2616 | 1.09 | 1.6 | 2.69 |
| *Cavolinia inflexa* | Pteropoda | Euthecosomata | SRRXXXXXX | 14107853 | 20521 | 1729 | 35.51 | 2.17 | 37.68 |
| *Clio cuspidata* | Pteropoda | Euthecosomata | SRRXXXXXX | 15473224 | 21307 | 1222 | 56 | 1.05 | 57.05 |
| *Clio pyramidata* | Pteropoda | Euthecosomata | SRRXXXXXX | 15596539 | 24705 | 1665 | 39.3 | 2.12 | 41.42 |
| *Clione antarctica* | Pteropoda | Gymnosomata | SRR1505107 | 20282761 | 19813 | 1915 | 29.53 | 2.62 | 32.16 |
| *Pneumodermopsis* sp. | Pteropoda | Gymnosomata | SRRXXXXXX | 18094803 | 18984 | 1396 | 51.35 | 1.24 | 52.59 |
| *Creseis acicula* | Pteropoda | Euthecosomata | SRRXXXXXX | 12027779 | 27003 | 1806 | 33.19 | 2.82 | 36.01 |
| *Creseis virgula* | Pteropoda | Euthecosomata | SRRXXXXXX | 14105581 | 26726 | 1824 | 34.12 | 3.06 | 37.18 |
| *Cuvierina atlantica* | Pteropoda | Euthecosomata | SRRXXXXXX | 28558114 | 20994 | 1966 | 28.34 | 2.99 | 31.33 |
| *Cymbulia sibogae* | Pteropoda | Pseudothecosomata | SRRXXXXXX | 13430464 | 14494 | 1166 | 57.8 | 1.83 | 59.63 |
| *Diacavolinia longirostris* | Pteropoda | Euthecosomata | SRRXXXXXX | 14469630 | 21220 | 1582 | 44.76 | 2.22 | 46.98 |
| *Diacria danae* | Pteropoda | Euthecosomata | SRRXXXXXX | 25218341 | 36533 | 1673 | 38.09 | 1.23 | 39.32 |
| *Diacria trispinosa* | Pteropoda | Euthecosomata | SRRXXXXXX | 13123731 | 16800 | 682 | 76.48 | 0.87 | 77.36 |
| *Haminoea antillarum* | Cephalaspidea | Outgroup | SRR1505111 | - | 39489 | 1821 | 35.68 | 5.68 | 41.36 |
| *Heliconoides inflatus* (ATL) | Pteropoda | Euthecosomata | SRRXXXXXX | 33421222 | 20283 | 1109 | 66.97 | 2.08 | 69.05 |
| *Heliconoides inflatus* (S ATL) | Pteropoda | Euthecosomata | SRRXXXXXX | 18330623 | 33123 | 1925 | 29.03 | 2.73 | 31.76 |
| *Heliconoides inflatus* (MED) | Pteropoda | Euthecosomata | PRJNA312154 | - | 25016 | 2434 | 7.83 | 2.2 | 10.03 |
| *Hyalocylis striata* | Pteropoda | Euthecosomata | SRRXXXXXX | 20300621 | 33352 | 2360 | 10.42 | 2.23 | 12.65 |
| *Limacina bulimoides* | Pteropoda | Euthecosomata | SRRXXXXXX | 18046287 | 28764 | 1799 | 34.54 | 2.81 | 37.35 |
| *Limacina antarctica* | Pteropoda | Euthecosomata | SRRXXXXXX | 19486196 | 28954 | 2221 | 15.92 | 2.07 | 17.99 |
| *Limacina lesueuri* | Pteropoda | Euthecosomata | SRRXXXXXX | 14648179 | 24430 | 1793 | 33.53 | 2.1 | 35.64 |
| *Limacina retroversa* | Pteropoda | Euthecosomata | PRJNA260534 | - | 40077 | 2376 | 10.62 | 2.53 | 13.15 |
| *Limacina trochiformis* | Pteropoda | Euthecosomata | SRRXXXXXX | 14412701 | 35829 | 2004 | 25.99 | 2.75 | 28.74 |
| *Peracle reticulata* | Pteropoda | Pseudothecosomata | SRRXXXXXX | 17876973 | 27998 | 1776 | 35.18 | 2.43 | 37.61 |
| *Philine angasi* | Cephalaspidea | Outgroup | SRR1505129 | 11881480 | 33219 | 2037 | 23.05 | 5.81 | 28.87 |
| *Pneumoderma violaceum* | Pteropoda | Gymnosomata | SRRXXXXX | 16362957 | 24989 | 2148 | 17.78 | 2.21 | 19.99 |
| *Spongiobranchaea australis* | Pteropoda | Gymnosomata | SRRXXXXX | 18191206 | 20998 | 1621 | 40.7 | 1.66 | 42.36 |
| *Styliola subula* | Pteropoda | Euthecosomata | SRRXXXXX | 16115592 | 29755 | 2062 | 20.76 | 2.33 | 23.09 |

**Table S3.**
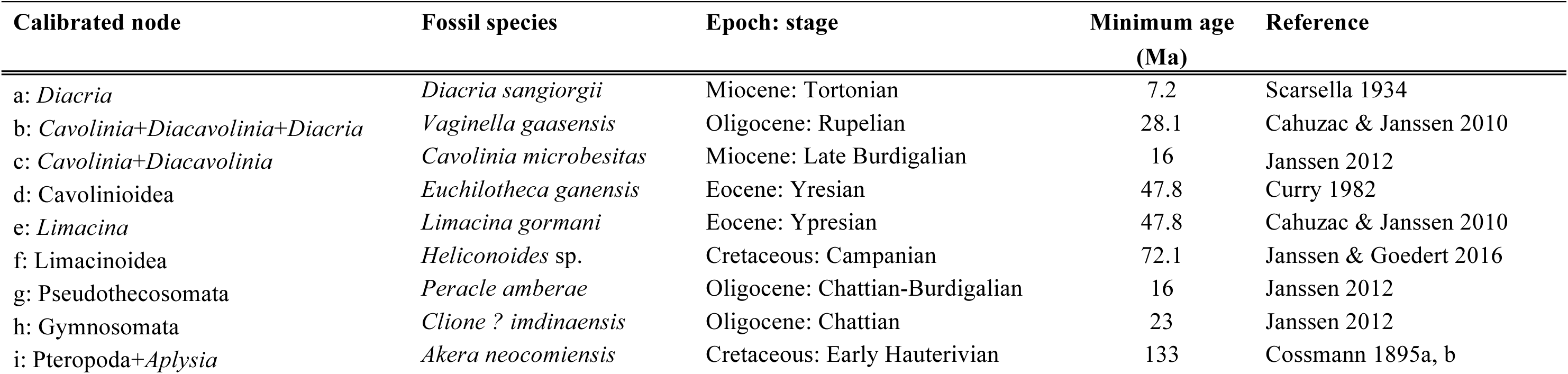
Nine fossil calibrations used in the molecular clock analyses. Letters refer to nodes labelled a-i in the chronogram of Figure 2.

| Calibrated node | Fossil species | Epoch: stage | Minimum age<br>(Ma) | Reference |
| --- | --- | --- | --- | --- |
| a: *Diacria* | *Diacria sangiorgii* | Miocene: Tortonian | 7.2 | Scarsella 1934 |
| b: *Cavolinia*+*Diacavolinia*+*Diacria* | *Vaginella gaasensis* | Oligocene: Rupelian | 28.1 | Cahuzac & Janssen 2010 |
| c: *Cavolinia*+*Diacavolinia* | *Cavolinia microbesitas* | Miocene: Late Burdigalian | 16 | Janssen 2012 |
| d: Cavolinioidea | *Euchilotheca ganensis* | Eocene: Yresian | 47.8 | Curry 1982 |
| e: *Limacina* | *Limacina gormani* | Eocene: Ypresian | 47.8 | Cahuzac & Janssen 2010 |
| f: Limacinoidea | *Heliconoides* sp. | Cretaceous: Campanian | 72.1 | Janssen & Goedert 2016 |
| g: Pseudothecosomata | *Peracle amberae* | Oligocene: Chattian-Burdigalian | 16 | Janssen 2012 |
| h: Gymnosomata | *Clione ? imdinaensis* | Oligocene: Chattian | 23 | Janssen 2012 |
| i: Pteropoda+*Aplysia* | *Akera neocomiensis* | Cretaceous: Early Hauterivian | 133 | Cossmann 1895a, b |

